# Brain-handedness associations depend on how and when handedness is measured

**DOI:** 10.1101/2024.02.15.580563

**Authors:** Link Tejavibulya, Corey Horien, Carolyn Fredricks, Bronte Ficek-Tani, Margaret L. Westwater, Dustin Scheinost

## Abstract

Hand preference is ubiquitous, intuitive, and often simplified to right- or left-handed. Accordingly, differences between right- and left-handed individuals in the brain have been established. Nevertheless, considering handedness as a binarized construct fails to capture the variability of brain-handedness associations across different domains or activities. Further, many cultures, environments, and generations impose right-handed norms, and handedness preferences can change over the lifespan. As a result, brain-handedness associations may depend on *how* and *when* handedness is measured. We used two large datasets, the Human Connectome Project-Development (HCP-D; n=465; age=5-21 years) and Human Connectome Project-Aging (HCP-A; n=368; age=36-100 years), to explore handedness preferences and brain-handedness associations. Nine items from the Edinburgh Handedness Inventory were associated with resting-state functional connectomes. We show that brain-handedness associations differed across the two cohorts. Moreover, these differences depended on the way handedness was measured. Given that brain-handedness associations differ across handedness measures and datasets, we caution against a one-size-fits-all approach to neuroimaging studies of this complex trait.

## Introduction

A preference for using one hand over another—or handedness—is a complex trait with well-studied neural correlates. Differences in the language^1–5^, motor^6–8^, and somatosensory^9–11^ networks have been consistently reported when comparing right- and left-handed individuals. For example, left-handed individuals exhibit more bilateral activation of brain regions during language tasks, suggesting a greater reliance on the right hemisphere for language processing^5^. The corpus callosum, which connects the two hemispheres, may be larger in left-handed individuals, indicating increased interhemispheric communication^12,13^. Recent work further suggests that differences related to handedness extend beyond localized regions to widespread functional connectivity differences, potentially affecting every canonical brain network^14–16^.

Nevertheless, how handedness preferences and accompanying brain-handedness associations form remains unclear. While individual differences in handedness are partly attributable to genetic factors^17,18^, environmental factors play an essential role in shaping the expression of handedness ^19–21^. For example, Western cultural preferences have historically encouraged right-hand dominance^22^. In some instances, young children were mandated to use their right hand for writing, using cutlery, and performing everyday tasks despite their natural preferences^23–25^. Moreover, although left-handed individuals comprise ∼10% of the population^22^, evidence suggests that rates of left-handedness have increased in recent years, perhaps due to growing tolerance of variation in this trait. A related challenge concerns the definition of handedness. Although it is often considered a binarized construct, such a definition fails to capture the variability of handedness preferences across domains or activities^26^. One may prefer to write with their right hand but throw a ball with their left. This heterogeneity is captured by validated handedness questionnaires, such as the Edinburgh^27–29^, the Annett^30^, and many others^31–33^. These generate ordinal or dimensional sum scores from items that span the full breadth of handedness use. Despite this rigor, the test-retest reliability of handedness scores changes over time^34^, suggesting the environment’s role in shaping hand preferences. As a result, brain-handedness associations may depend on *how* and *when* handedness is measured.

In the present study, we used two large datasets, the Human Connectome Project-Development (HCP-D; n = 465; age = 5-21 years) and Human Connectome Project-Aging (HCP-A; n = 368; age = 36-100 years), to explore handedness preferences and brain-handedness associations. We hypothesized that brain-handedness associations would differ across these datasets for multiple different measures of handedness. First, we investigated group differences between pairs of granular measures of handedness on a behavioral level. We then investigated group differences in brain-handedness associations. Finally, we compared how brain-handedness associations differed for every granular measure of handedness between cohorts. Our findings show that, although the covariance among granular handedness measures was similar between the datasets, patterns of brain-handedness associations differed significantly. These results suggest that developmental^35–37^, environmental^19–21 34^, and generational^26,38,39^ factors shape brain-handedness associations. Given that brain-handedness associations differ across handedness measures and datasets, we caution against a one-size-fits-all approach to neuroimaging studies of this complex trait.

## Results

### Behavioral correlations

We analyzed behavioral and neuroimaging data from the Human Connectome Project (HCP) Lifespan Studies. The Human Connectome Project-Development (HCP-D; brain and behavior n = 465, behavior only n= 652) was used as the younger cohort. The Human Connectome Project-Aging (HCP-A; brain and behavior n = 368, behavior only n = 724) was used as the older cohort. The HCP-D consisted of individuals aged 5 to 21 years, whereas the HCP-A included participants between ages 36 and 100 years. Measures of handedness were determined from individuals’ self-reported preferences for their dominant hand using nine items from the Edinburgh Handedness Inventory^40^. The following nine items were assessed: writing, throwing, scissors-use, toothbrush-use, using a knife without a fork, spoon-use, sweeping (using a broom), lighting a match, and opening a box. Individuals responded using a 5-point Likert scale, ranging from 1 = *Always left* to 5 *= Always right*, to indicate their hand preference for each item.

When assessing the covariance of individual Edinburgh Handedness Inventory (EHI) items, we found that all items were significantly correlated for both the HCP-D (Fig. 1A) and the HCP-A cohorts (Fig. 1B). Patterns of Spearman correlations (ρ) were similar between the two generations (r=0.85 between correlation matrices). Nevertheless, correlations were significantly stronger in the HCP-A cohort (Table S2) than among the younger HCP-D cohort (Table S1; Fig. S1), these differences were further validated by performing significance tests between pairs of measures of handedness (Table S3, S4). These results suggest that the EHI measures captured similar information for each cohort.

**Fig. 1:**
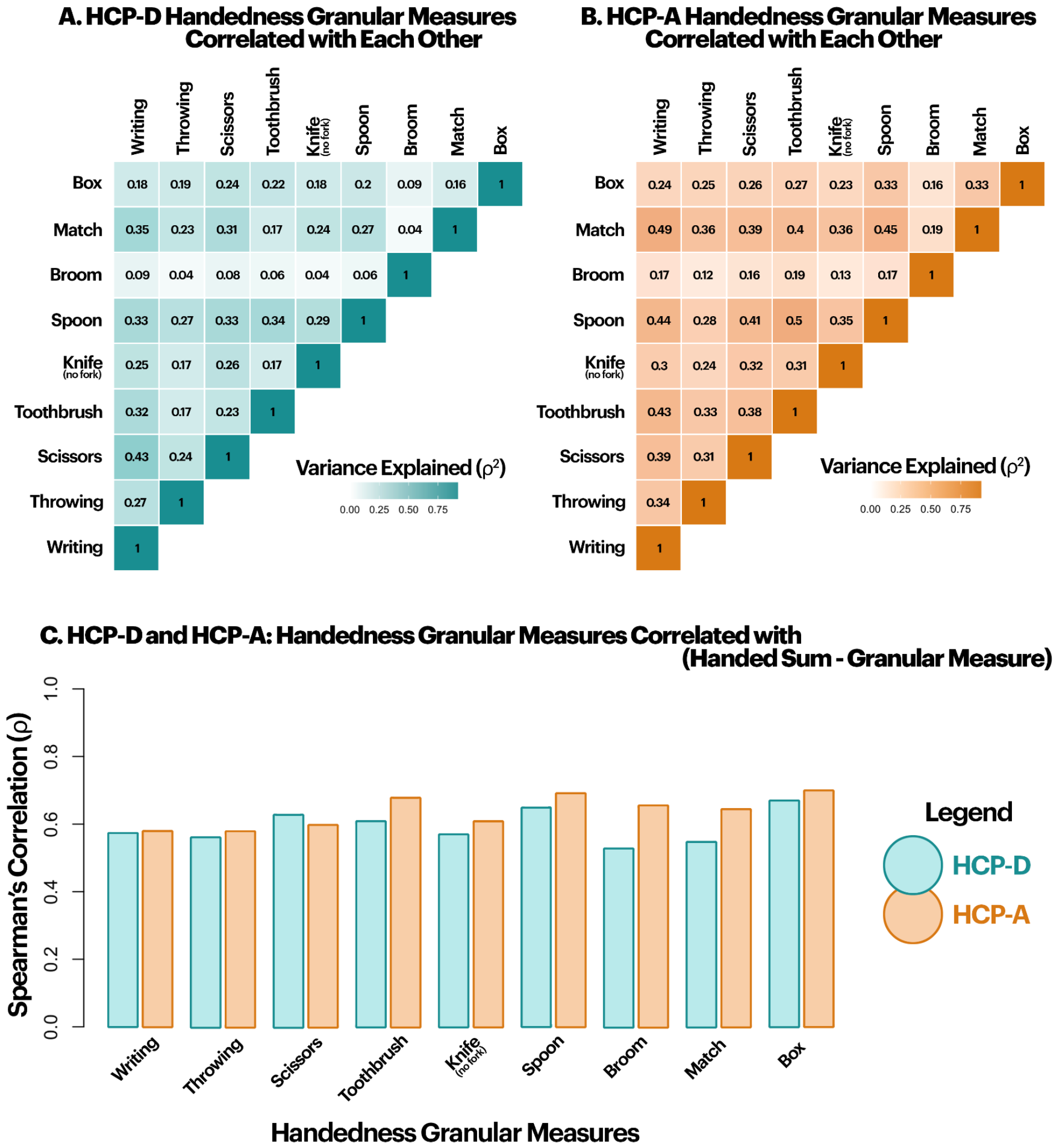
Behavioral correlations between pairs of granular measures of handedness and the sum score. (A) Spearman’s variance explained between pairs of granular measures of handedness for the HCP-D. (B) Spearman’s variance explained between pairs of granular measures of handedness for the HCP-A. (C) Granular measures of handedness correlated with the handedness sum score minus that particular granular measure. Spearman’s correlations (ρ) for the HCP-D are displayed in blue and HCP-A in orange.

As the current gold standard for measuring handedness utilizes a sum score of EHI items, next, we investigated how well a given item performed relative to the sum score.

Spearman’s ρ’s were calculated between each granular measure and the handedness sum score, minus that particular granular measure, separately for the two cohorts (Fig. 1C). In general, correlations between handedness sum score and any given item were stronger for the HCP-A dataset than for the HCP-D cohort.

### Brain-handedness associations

We first generated 268-node functional connectomes for each participant to investigate brain-behavior associations using resting-state fMRI data. Then, we assessed edge-wise Spearman’s correlations with EHI sum scores and individual items separately for the HCP-D and HCP-A datasets. We used the Network-Based Statistics (NBS)^41^ to identify components where functional connectivity was significantly associated with handedness while controlling for the family-wise error rate. As thresholding results based on significance can modulate observed group differences^42^, we also examined the effect sizes for handedness separately for the HCP-D and the HCP-A datasets. Out of the 35,778 total edges in a connectome, 5.5% and 6.27% were identified as statistically (p<0.05, corrected) different for right- and left-handed groups in the HCP-D cohort, respectively. In the HCP-A cohort, 4.39% and 4.96% of edges were significant (p<0.05, corrected) for right- and left-handed groups, respectively. In each group, differences were widespread and complex, with contributions from every node and canonical brain network (Fig. 2). Next, we explored differences between right-handed males and females to benchmark meaningful handedness differences. The neural correlates of self-reported sex are widely studied^43–46^ and often controlled for in fMRI studies^47,48^. The effect size distribution for handedness was wider than for sex for both datasets, indicating that handedness generally exhibited larger effect sizes than sex.

**Fig. 2:**
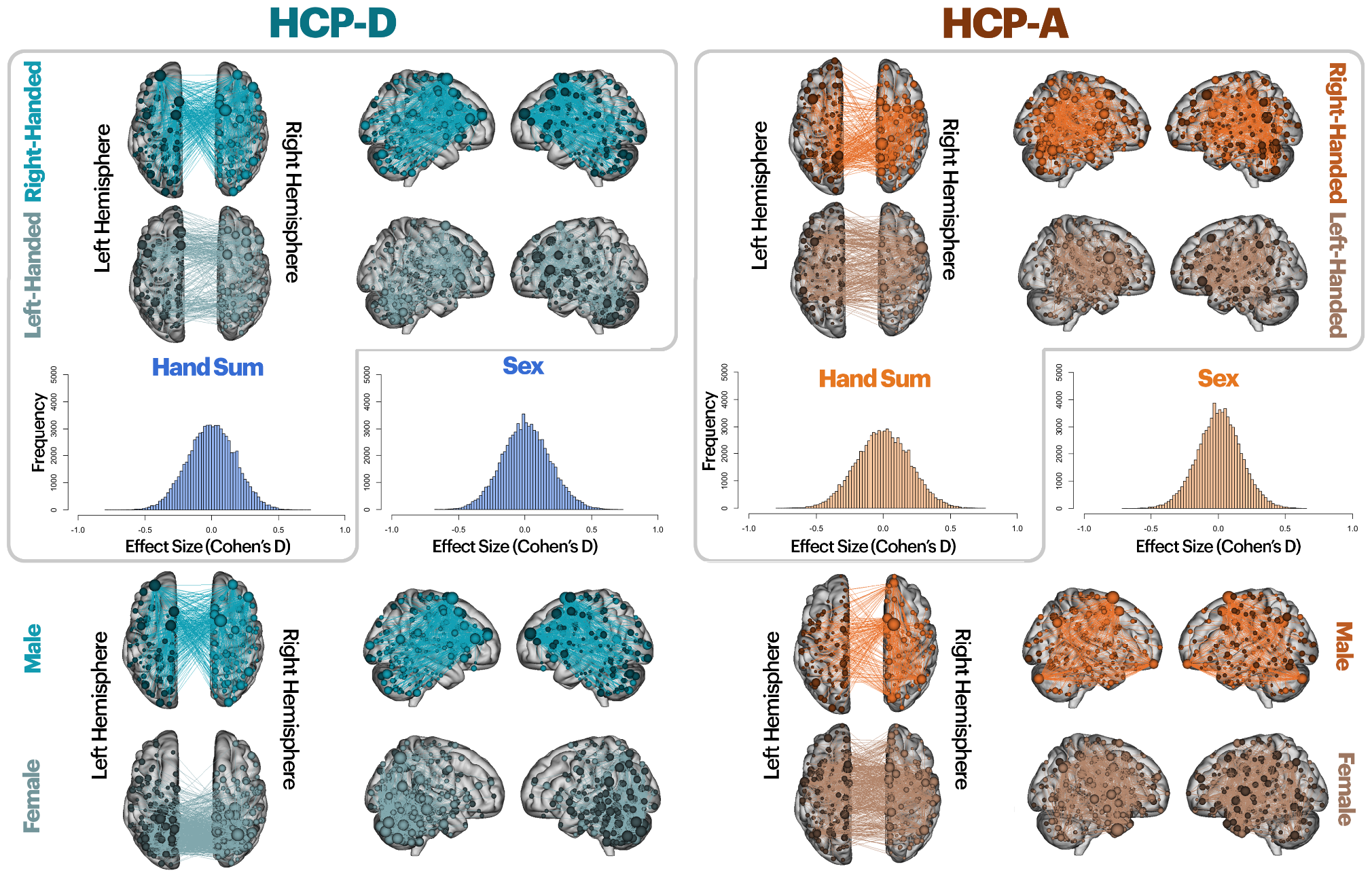
Handedness vs Sex: Boxed: Edges significantly different between left- and right-handed groups were identified based on the handedness sum score. Effect sizes were then calculated for all 35,778 edges and shown on a histogram. This procedure was performed separately for the two cohorts with the HCP-D (on the left in blue) and the HCP-A (on the right in orange). Unboxed: Edges significantly different between right-handed males and females were identified based on self-reported sex. Effect sizes were then calculated for all 35,778 edges and shown on a histogram. This procedure was performed separately for the two cohorts with the HCP-D (on the left in blue) and the HCP-A (on the right in orange). Thresholds for 3D brain visualizations (thresholded at a minimum number of significant edges connected to a specific node) were as follows: [HCP-D] right-handed: 35, left-handed: 35, male: 80, female: 75 [HCP-A] right-handed: 25, left-handed, 25, male: 60, female: 50.

### Exploring effect sizes in granular measures

Next, we examined how effect sizes differed for individual EHI items. Effect sizes were calculated separately for the HCP-D and the HCP-A datasets. This procedure resulted in a matrix where each element represented the Cohen’s D between an edge’s functional connectivity and hand preference for each EHI item and dataset. Within a dataset, effect sizes for all individual handedness items showed similar distributions, except for opening a box and broom use. (Fig. S2). The box and broom-use items had smaller effect sizes than the handedness sum score overall. Komolgorov-Smirnov test statistics were performed to compare the effect size distribution between the two cohorts (Table S5). All distributions were significantly different, with the HCP-A generally being wider (i.e., larger effects are observed in the HCP-A).

### Comparing brain-handedness associations between cohorts

We formally compared the brain-handedness associations between cohorts, using edge-wise z-tests (Fig. 3). Widespread differences in brain-handedness associations between the HCP-D and the HCP-A cohorts for the handedness sum score were observed. These differences account for 12.5% of edges (out of 35,778). While results were widespread across the brain, a large cluster of edges was located in posterior regions. Differences were also calculated between left- and right-handed groups based on individual handedness items (Fig. 3: periphery). While measures such as throwing, scissors-use, knife-use, writing, and match-use had significant edges that were comparable to or higher than that of the handedness sum score (10.88%, 11.06%, 12.79%, 10.2%, and 14.2%, respectively), not all measures demonstrated as large of differences between the two cohorts. As expected based on our earlier results (Fig. S2), box opening and broom use were associated with a lower percentage of significant edges at 6.31% and 3.87%, respectively. When we explored the percentage of overlap between each handedness item and the handedness sum score, these values ranged from 1.24% (broom-use) to 7.18% (writing), indicating that the functional correlates of a given item differ across the cohorts. In other words, while differences in brain-handedness associations between the cohorts exhibit widespread patterns, each granular measure still has distinct differences.

**Fig. 3:**
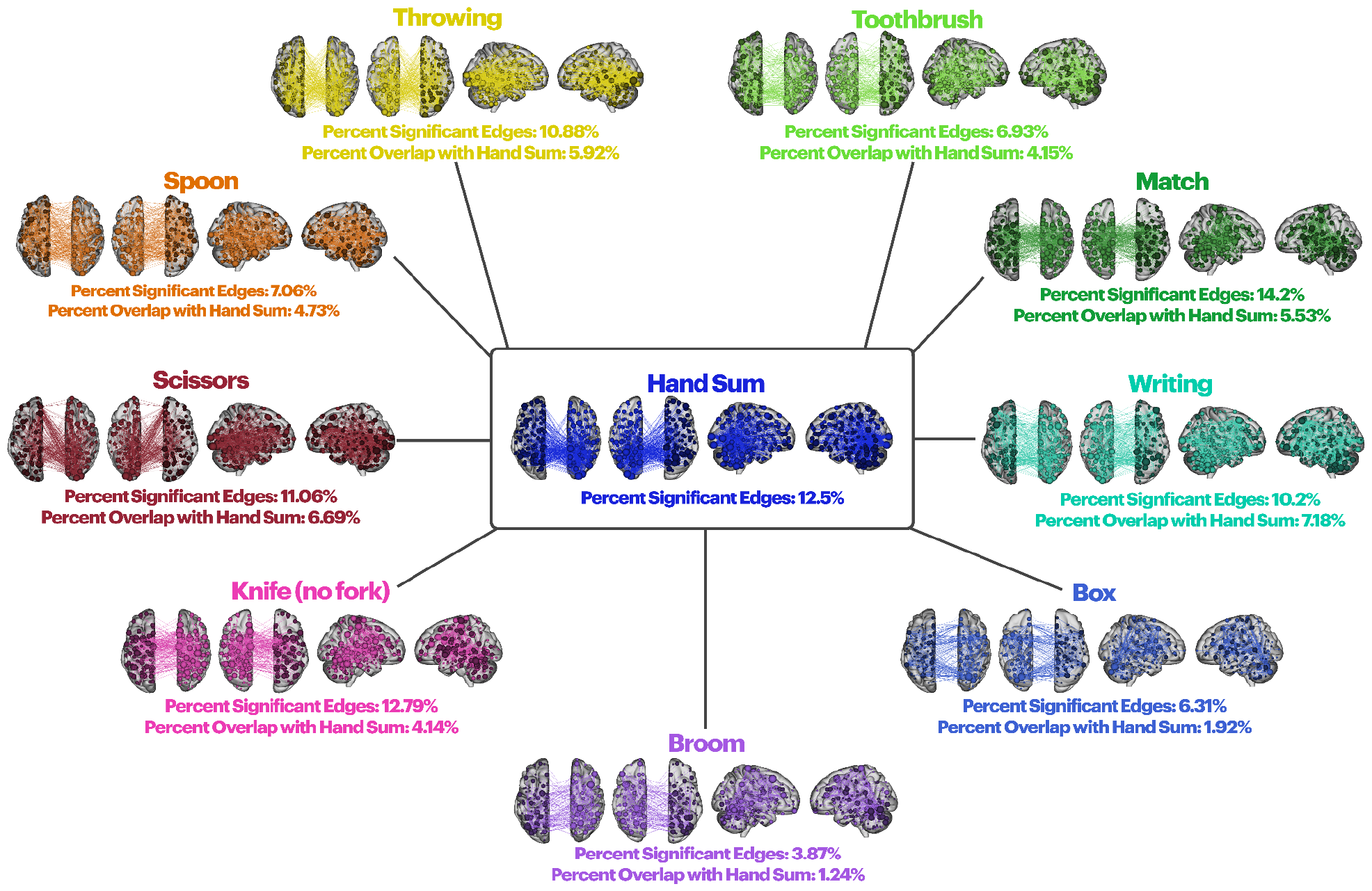
Edges significantly different between the two cohorts. Edges exhibited significant differences in brain-handedness associations between the HCP-D and the HCP-A. Visualizations were thresholded (minimum number of edges connected to a specific node) for interpretability at 35 for spoon-use, 55 for throwing, 35 for toothbrush-use, 70 for match-use, 50 for writing, 35 for box opening, 25 for broom-use, 70 for knife-use (no fork), and 55 for scissors-use. The percentage of significant edges for each granular measure (p < 0.05) and the percentage of overlapping edges with the handedness sum score are displayed below each 3D brain plot. For the percent of overlap with the handedness sum score, all edges that yielded significance for both the granular measure and the sum score were summed up and calculated as a percentage of the whole connectome.

### Examining covariance of brain-handedness effect sizes

To explore how similar the brain-handedness associations were across EHI items, we correlated effect size maps (Fig. S2) for all pairwise item combinations (Fig. 4). In HCP-D, effect size maps were weakly correlated or uncorrelated, except for toothbrush-use, scissors-use, throwing, and writing. In contrast, all item-level effect size maps were strongly correlated in HCP-A. In other words, for HCP-A, edges that show a strong correlation with one EHI item tend to show strong correlations with other EHI items. For HCP-A, these correlation patterns (Fig. 4B) resemble the behavioral variance explained (Fig. 1B). In contrast, HCP-D does not show this pattern. These results were further validated by significance tests between pairs of correlations (Table S6, S7).

**Fig. 4:**
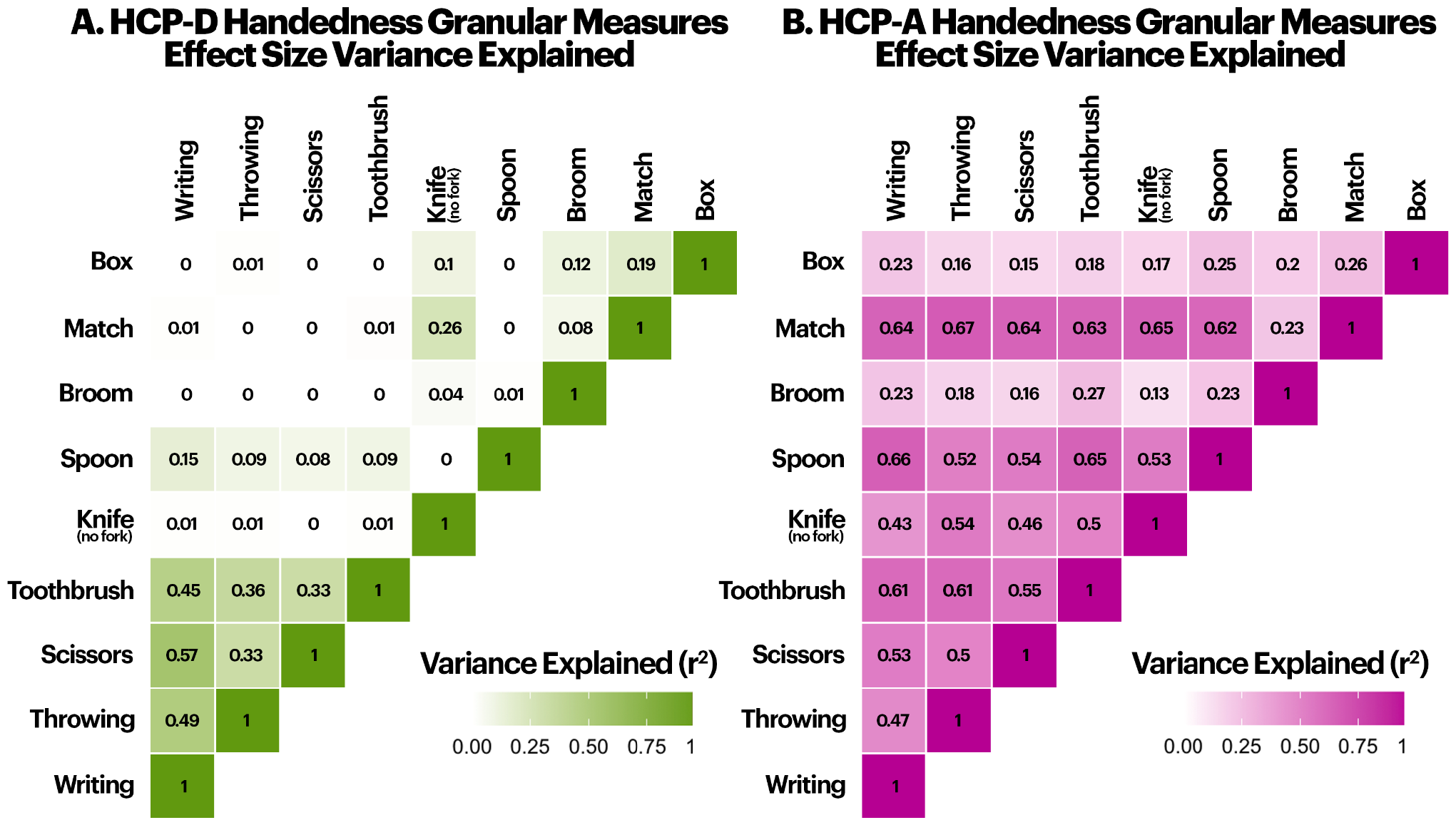
Effect size variance explained for all granular measures of handedness calculated separately for the two cohorts. Correlations between brain-handedness effect size maps for pairs of specific granular measures shown for HCP-D (left) and HCP-A (right).

To relate brain-handedness associations (Fig. 4A and 4B) and behavioral explained variances (Fig. 1A and 1B) between pairwise EHI items, behavioral and effect size correlations were visualized (Fig. 5) and subsequently correlated. As anticipated, brain-behavior correlations were weaker in the HCP-D (r=0.29) than in the HCP-A (r=0.81) dataset (z=3.41, p-val<0.01).

**Fig. 5:**
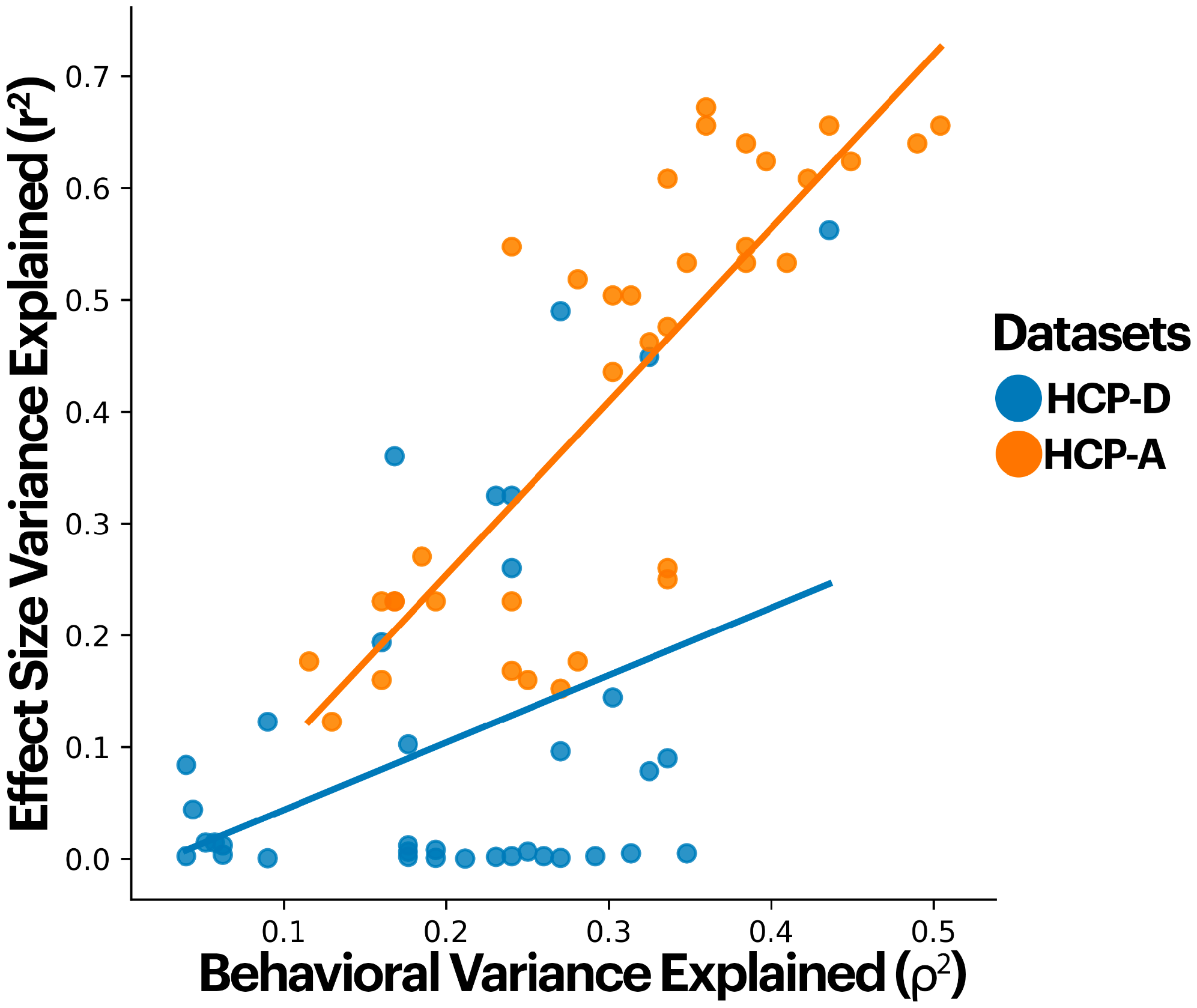
Correlations plots between brain and behavioral variance explained by cohort. Scatter plot of effect size variance explained (r^2^) and behavioral variance explained (ρ^2^) as seen from figures 1 and 4, respectively.

These results suggest that—although, behaviorally, younger cohorts may have similar covariance between different items—the underlying functional organization of the brain for each item appears distinct.

## Discussion

This study investigated brain-handedness associations across multiple granular measures of handedness and how these associations differ across age groups. Importantly, brain-handedness associations differed across the two cohorts. Further, these differences depended on the way handedness was measured. Our results, first and foremost, suggest that the question “Are you right-or left-handed?” may be oversimplified for brain-handedness associations. Brain-handedness associations likely depend on how and when handedness is measured in the lifespan.

Differences in brain-handedness associations between the HCP-D and HCP-A may be developmental^35–37^, environmental^19–21 34^, or generational^26,38,39^. Nevertheless, pinpointing the exact mechanisms underlying our results is difficult. A natural interpretation is that these associations crystalize over the lifespan and, thus, are stronger in the HCP-A. While differences between the two cohorts point to a developmental effect, data from the HCP-D and HCP-A are not longitudinal. We cannot ascertain the full extent of how developmental trajectories influence handedness preferences and their underlying neural correlates. Thus, it is difficult to conclude whether the observed differences reflect a developmental effect that has changed over time.

The environment may also play a role in the observed associations. Specific measures, such as writing and throwing, are taught in schools from a very young age. Yet others, like knife-use, broom-use, and lighting a match, would be less common behaviors in a younger cohort.

Therefore, brain-handedness associations for certain measures may not be particularly informative in a younger cohort due to lower engagement rates. Conversely, scissors-use correlations are marginally higher for HCP-D than HCP-A, which may be because younger cohorts are more actively using scissors in classroom activities. Additionally, we cannot account for all individual differences of participants. For instance, athletes train to utilize their hands or perform motor movements with more nuance than the general population. Training and using hands for specific tasks will moderate brain-handedness associations.

Cultural preferences are another environmental factor that shapes brain-handedness associations. Data from the Human Connectome Projects were collected within the United States. While cultural differences in handedness are crucial^49,50^, we were limited in assessing these effects as all participants came from a Western, educated, industrialized, rich, and democratic (WEIRD) population^51,52^. In contrast, Eastern cultures have been known to impose right-handedness on children from a very young age^19,20^, thus further affecting how individuals from these cultures exhibit handedness preference changes later on^53^. Studies surrounding cultural changes offer a unique opportunity to more clearly disentangle the interactions between genetics, epigenetics, and the environment. Nonetheless, for these studies to be successfully conducted, we would require data to be collected from many different countries and cultures.

In addition to environmental and age effects, generational effects in handedness exist.

Generational effects can be broadly defined as an age-by-environment interaction where individuals of different ages live in different environments. Older individuals may have been more adept at using their right hands overall since older generations were more likely forced to use their right hands. The percentage of left-handed individuals has increased as these cultural influences change. Similarly, handwriting is less common in younger generations as digital communication increases. Because the datasets used for these analyses come from different generations, we can not directly interpret differences between the cohorts as “changes over time”. Unfortunately, the HCP-D and HCP-A consist of wide age ranges. We could not define generational groups (i.e., Millennials) stringently. Additionally, given the limited number of left-handed individuals at any particular age, we were not powered to split the cohorts into narrower age bins. Results may indicate a generational effect (i.e., older generations were encouraged to be right-hand dominant regardless of preference), but testing this hypothesis in these data is difficult due to confounding effects of age.

Longitudinal studies over different generations and countries are likely needed to disentangle our results’ mechanisms. While it is tempting to infer that brain-handedness associations for an individual do not change over time, the test-retest reliability for handedness measures is imperfect^54^. Handedness preferences may change over time^55^. The extent to which these changes occur also depends on the developmental periods during observed time points. Thus, it is essential to consider that EHI scores reflect an individual at a given time, and it can not be concluded that results will remain the same for future time points. Nevertheless, collecting data over decades is a difficult task. Similarly, animal studies offer the possibility of longitudinal studies across lifespans, and greater experimental manipulations exist. However, animals do not have the same motor capacities as humans and cannot self-report handedness preferences beyond pure observations. Thus, studies of brain-handedness associations in animal models cannot address these questions. Longitudinal datasets (such as the Adolescent Brain Cognitive Development Study) may represent a starting point for further investigations.

In conclusion, we examine brain-handedness associations from multiple measures across an extensive age range. We demonstrated that not all handedness measures are equal, and each measure exhibits associations and cohort differences. These handedness differences could be attributed to many factors, including generational, cultural, and individual differences. Regardless of mechanisms, brain-handedness associations likely depend on how and when handedness is measured.

## Methods

### Datasets: Human Connectome Project-Development (HCP-D) & Human Connectome Project-Aging (HCP-A)

Two datasets from the Human Connectome Project, Development^56–58^ and Aging^57,59^, were used throughout this study to represent the younger and older cohorts, respectively. Data were obtained from two separate populations. Table 1 summarizes each dataset’s participant population, scanner information, and exclusion criteria.

**Table 1:** Dataset description.

| Table 1: Dataset description |  |  |
| --- | --- | --- |
| Dataset | Human Connectome Project-Development<sup>56–58</sup><br>(HCP-D) | Human Connectome Project-Aging<sup>57,59</sup><br>(HCP-A) |
| Age Range | 5-22 years | 36-100 years |
| Number of participants | Behavior only: 652<br>Brain and behavior: 465 | Behavior only: 724<br>Brain and behavior: 368 |
| fMRI scans | Resting-state | Resting-state |
| Length of scan | 10 mins | 10 mins |
| Scanner | 3T Siemens scanners | 3T Siemens scanners |
| Subject exclusion criteria | Missing behavioral/brain data<br>Excessive motion (>0.1mm) | Missing behavioral/brain data<br>Excessive motion (>0.1mm) |

### Behavioral Measures of Handedness

For handedness measures, participants completed a subset of the Edinburgh Handedness Inventory (EHI) items where they were asked which hand they preferred to write, throw a ball, use a spoon, and brush their teeth, upper hand when using a broom, light a match, box, use scissors and use a knife (without fork) with. Participants rated each item on a Likert scale of Always left, Sometimes left, Indifferent, Sometimes right, and Always right. These granular measures were used to classify the extent to which subjects were left-or right-sided.

Our handedness sum score was simply a summation of all nine values for each subject to determine overall handedness (unless specified otherwise).

### Preprocessing and generating connectomes

The HCP-D and HCP-A datasets were analyzed with identical processing pipelines. Structural scans were first skull stripped using an optimized version of the FMRIB’s Software Library (FSL)^60^ pipeline^61^. Functional images were motion-corrected using SPM12^62^. All further analyses were performed using BioImage Suite^63^. Several covariates of no interest were regressed from the data, including linear and quadratic drifts, mean cerebral-spinal-fluid (CSF) signal, mean white-matter signal, and mean gray matter signal. For additional control of possible motion-related confounds, a 24-parameter motion model (including six rigid-body motion parameters, six temporal derivatives, and these terms squared) was regressed from the data.

The data were temporally smoothed with a Gaussian filter (approximate cutoff frequency=0.12 Hz).

Nodes were defined using the Shen 268-node brain atlas^64^, which includes the cortex, subcortex, and cerebellum as described in prior work. The atlas was warped from MNI space into single-subject space via a series of linear and non-linear transformations calculated using a previously validated algorithm^65^, implemented in BioImage Suite. This involved computation of the mean time courses for each of the 268 nodes (i.e., averaging the time courses of all constituent voxels). Node-by-node pairwise correlations were computed, and Pearson correlation coefficients were Fisher z-transformed to yield symmetric 268×268 connectivity matrices, in which each matrix element represents the connectivity strength between two individual nodes (i.e., ‘edge’).

### Correlations for behavioral measures

Pairwise correlations for handedness items were calculated using Spearman’s correlation (ρ). These values were then plotted on a heat map to demonstrate how similar handedness granular measures were to others.

### Comparing correlation significance between datasets

Correlation coefficients were converted to Z-scores, and p-values were subsequently calculated.

### Identifying significant edges and nodes between handedness and sex

Differences between the left-handed and right-handed groups and right-handed males and right-handed females were estimated using Network-Based Statistics (NBS)^41^ (component-determining threshold Z=1.96, 2-tailed, K=5000 permutations). Age was additionally used as a covariate to control for these differences. NBS is analogous to cluster-based correction and solves the statistical problem of multiple comparisons in a whole-brain connectivity analysis. In NBS, using the difference between groups, the largest fully connected network of suprathreshold edges, or “component,” is identified, and its extent is defined as the number of edges it comprises. We plotted edges and nodes on two visualizing modalities to visualize these results: ball-and-stick figures on 3D brains. Due to the complicated nature of our results and the high number of edges in a connectome, some results have had to be thresholded for interpretability. Thresholds only visualize edges connected to nodes with a minimum number of connections.

### Whole brain effect sizes

We separately quantified the effect size of connectivity differences between left- and right-handed individuals for all edges using Cohen’s D for the HCP-D and HCP-A. To help put these whole-brain differences into a comparable context, we compared our effect sizes relative to sex differences, which have been known to have large effect sizes and are tightly controlled for in fMRI studies^47,48^. Effect sizes for sex were calculated using only primarily right-handed individuals relative to each granular measure. Cohen’s D was then calculated for the difference between male and female participants (based on self-reported sex).

### Differences between whole brain effect size

Cohen’s d’s were first converted to Pearson’s coefficients (r) and then subsequently z-scored to compare correlations between effect sizes between the HCP-D and the HCP-A cohorts. These z-scores were then thresholded at p-values of below 0.05 to visualize edges in a connectome that were deemed significantly different between the two cohorts and subsequently visualized on 3D brain plots using BioImage Suite.

### Correlations and variance explained for effect sizes

Correlations for each behavioral measure were calculated using Pearson’s r between each measure. These values were then squared to give variance explained (r^2^) and these r^2^ were subsequently plotted on a heat map to demonstrate how similar effect size maps for each handedness granular measure were to one another.

