## Supplementary Information and Tables for "Brain-handedness associations depend on how and when handedness is measured"

### Supplementary Figures

| | Writing | Throwing | Scissors | Toothbrush | Knife (no fork) | Spoon | Broom | Match | Box |
|---|---|---|---|---|---|---|---|---|---|
| Box | 0.03 | <0.01 | 0.20 | <0.01 | <0.01 | <0.01 | 0.11 | <0.01 | |
| Match | 0.01 | <0.01 | 0.16 | <0.01 | <0.01 | <0.01 | <0.01 | | |
| Broom | 0.02 | 0.02 | 0.02 | <0.01 | 0.01 | <0.01 | | | |
| Spoon | <0.01 | 0.13 | 0.05 | <0.01 | <0.01 | | | | |
| Knife (no fork) | 0.03 | <0.01 | 0.50 | <0.01 | | | | | |
| Toothbrush | 0.01 | <0.01 | 0.47 | | | | | | |
| Scissors | <0.01 | 0.02 | | | | | | | |
| Throwing | 0.16 | | | | | | | | |
| Writing | | | | | | | | | |

**Fig. S1:** p-values for comparing significance in behavioral correlations between the HCP-D (Fig. 1A) and the HCP-A (Fig. 1B).

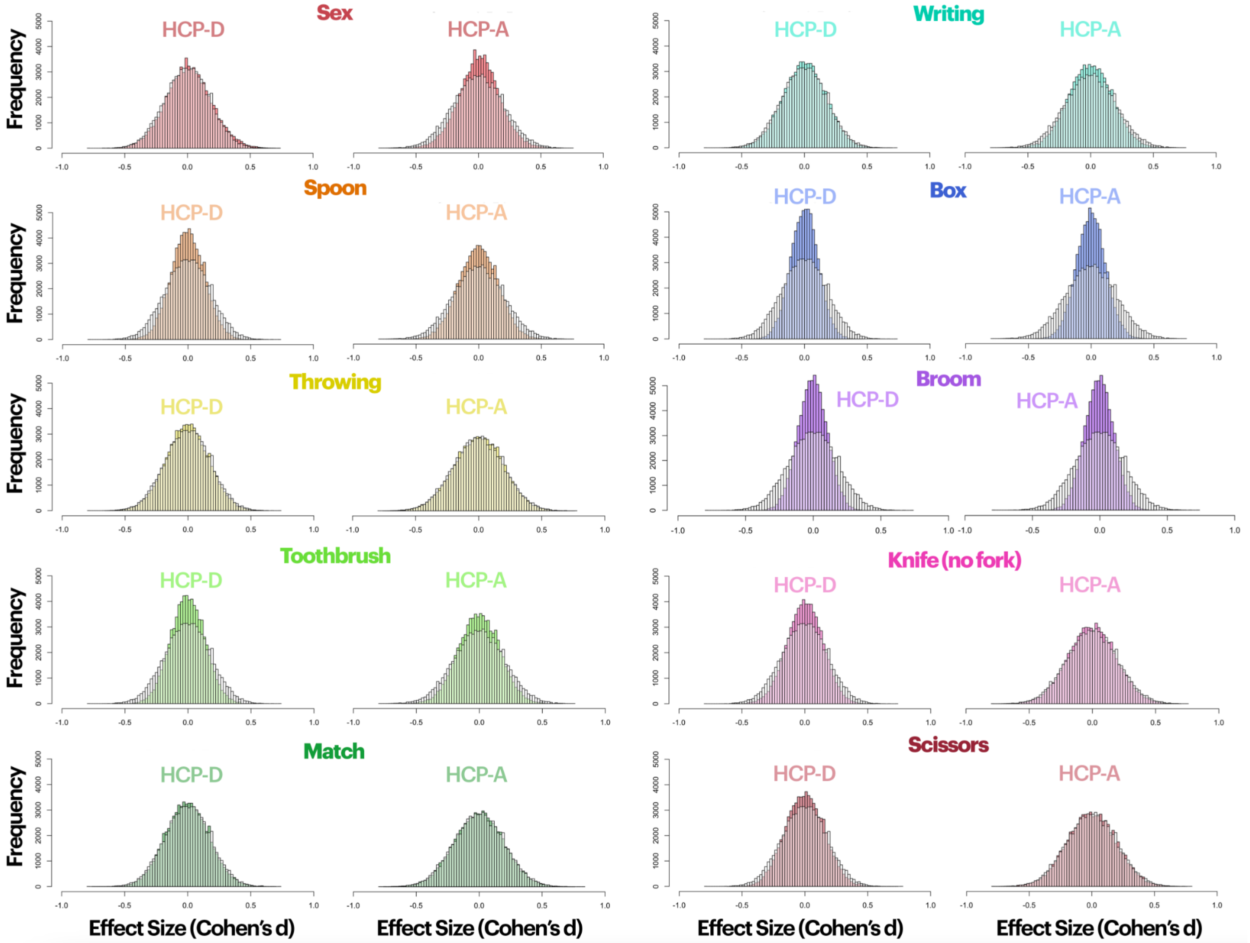

**Fig. S2:** Effect size differences between edges of a connectome for HCP-D compared to HCP-A for each granular measure of handedness (colored histograms) compared to effect sizes for handedness sum score (white bars). To show relevance to well-studied differences, such as sex, effect sizes for differences between right-handed males and females are shown on the top left panel.

#### Supplementary Tables

| | Extremely left (n) | Left (n) | Indifferent (n) | Right (n) | Extremely Right (n) |
|---|---|---|---|---|---|
| **Writing** | 66 | 1 | 0 | 18 | 567 |
| **Throwing** | 41 | 11 | 20 | 103 | 477 |
| **Scissors** | 45 | 8 | 22 | 61 | 516 |
| **Toothbrush** | 51 | 12 | 50 | 87 | 451 |
| **Knife (no fork)** | 29 | 18 | 28 | 73 | 340 |
| **Spoon** | 57 | 11 | 31 | 75 | 477 |
| **Broom** | 73 | 63 | 96 | 74 | 182 |
| **Match** | 26 | 10 | 24 | 57 | 371 |
| **Box** | 28 | 16 | 122 | 86 | 235 |

**Table S1:** Edinburgh Handedness Inventory item endorsement in HCP-D. Values indicate the number of participants endorsing each Likert value.

| | Extremely left (n) | Left (n) | Indifferent (n) | Right (n) | Extremely Right (n) |
|---|---|---|---|---|---|
| **Writing** | 81 | 3 | 0 | 18 | 623 |
| **Throwing** | 50 | 12 | 23 | 87 | 552 |
| **Scissors** | 45 | 9 | 20 | 71 | 580 |
| **Toothbrush** | 72 | 14 | 39 | 77 | 523 |
| **Knife (no fork)** | 55 | 16 | 32 | 84 | 538 |
| **Spoon** | 62 | 17 | 30 | 101 | 515 |
| **Broom** | 84 | 65 | 108 | 97 | 370 |
| **Match** | 59 | 11 | 18 | 72 | 564 |
| **Box** | 49 | 26 | 134 | 134 | 381 |

**Table S2:** Edinburgh Handedness Inventory item endorsement for HCP-A. Values indicate the number of participants endorsing each Likert value.

| Z scores | Writing | Throwing | Scissors | Toothbrush | Knife (no fork) | Spoon | Broom | Match | Box | Hand Sum |
|---|---|---|---|---|---|---|---|---|---|---|
| **Writing** | | | | | | | | | | |
| **Throwing** | 1.389 | | | | | | | | | |

| | | | | | | | | | |
|---|---|---|---|---|---|---|---|---|---|
| **Scissors** | -3.1728 | 2.2473 | | | | | | | |
| **Toothbrush** | 2.741 | 3.728 | 0.719 | | | | | | |
| **Knife (no fork)** | 2.176 | 4.124 | 0.676 | 4.075 | | | | | |
| **Spoon** | 3.667 | 1.512 | 1.954 | 5.049 | 3.223 | | | | |
| **Broom** | 2.247 | 2.301 | 2.414 | 3.303 | 2.542 | 3.258 | | | |
| **Match** | 2.773 | 4.344 | 1.401 | 5.640 | 4.963 | 4.124 | 4.178 | | |
| **Box** | 2.151 | 4.006 | 1.286 | 3.059 | 3.032 | 4.469 | 1.607 | 5.499 | |
| **Hand Sum** | 0.697 | 3.874 | -0.0979 | 6.111 | 3.269 | 5.392 | 2.679 | 6.543 | 3.190 |

**Table S3:** Table of z scores of behavioral correlation comparisons between pairs of granular measures.

| P values | **Writing** | **Throwing** | **Scissors** | **Toothbrush** | **Knife (no fork)** | **Spoon** | **Broom** | **Match** | **Box** | **Hand Sum** |
|---|---|---|---|---|---|---|---|---|---|---|
| **Writing** | | | | | | | | | | |
| **Throwing** | 0.165 | | | | | | | | | |
| **Scissors** | 0.00151 | 0.0246 | | | | | | | | |
| **Toothbrush** | 0.00612 | 0.00019 | 0.471 | | | | | | | |
| **Knife (no fork)** | 0.0296 | 3.72E-05 | 0.499 | 4.61E-05 | | | | | | |
| **Spoon** | 0.00024 | 0.131 | 0.0507 | 5.05E-07 | 0.00127 | | | | | |
| **Broom** | 0.02461 | 0.02139 | 0.01578 | 0.00096 | 0.0110 | 0.00112 | | | | |
| **Match** | 0.00555 | 1.40E-05 | 0.1611 | 1.70E-08 | 6.93E-07 | 3.73E-05 | 2.94E-05 | | | |
| **Box** | 0.03151 | 6.17E-05 | 0.1985 | 0.00222 | 0.00243 | 7.85E-06 | 0.10813 | 3.83E-08 | | |
| **Hand Sum** | 0.4861 | 0.00011 | 0.9212 | 9.89E-10 | 0.00108 | 6.95E-08 | 0.00739 | 6.04E-11 | 0.00142 | |

**Table S4:** Table of p values of behavioral correlation comparisons between pairs of granular measures.

| | **Komolgorov-Smirnov Statistic** | **P-Value** |
|---|---|---|
| **Hand Sum** | 2.90E-2 | 1.23E-26 |
| **Writing** | 1.28E-2 | 1.52E-05 |
| **Throwing** | 4.55E-2 | 4.41E-65 |
| **Scissors** | 6.00E-2 | 9.50E-113 |
| **Toothbrush** | 4.74E-2 | 1.73E-70 |
| **Knife (no fork)** | 7.50E-2 | 1.43E-175 |

| | | |
|---|---|---|
| **Spoon** | 4.57E-2 | 1.77E-65 |
| **Broom** | 2.54E-2 | 1.69E-20 |
| **Match** | 4.21E-2 | 1.20E-55 |
| **Box** | 1.67E-2 | 3.93E-09 |
| **Sex** | 3.44E-2 | 1.96E-37 |

**Table S5**: Kolmogorov-Smirnov tests were used for differences between the effect size distributions for HCP-D and HCP-A.

| Z scores | **Writing** | **Throwing** | **Scissors** | **Toothbrush** | **Knife (no fork)** | **Spoon** | **Broom** | **Match** | **Box** | **Hand Sum** |
|---|---|---|---|---|---|---|---|---|---|---|
| **Writing** | | | | | | | | | | |
| **Throwing** | -6.0607 | | | | | | | | | |
| **Scissors** | -10.294 | 42.819 | | | | | | | | |
| **Toothbrush** | 42.945 | 67.638 | 56.671 | | | | | | | |
| **Knife (no fork)** | 164.567 | 197.422 | 145.957 | 188.405 | | | | | | |
| **Spoon** | 138.306 | 112.299 | 124.359 | 154.846 | 182.568 | | | | | |
| **Broom** | 94.752 | 77.548 | 74.153 | 121.354 | 30.292 | 77.198 | | | | |
| **Match** | 221.044 | 225.133 | 195.61 | 220.735 | 105.311 | 197.751 | 41.526 | | | |
| **Box** | 91.164 | 63.032 | 68.005 | 86.694 | 20.290 | 100.098 | 22.748 | 16.447 | | |
| **Hand Sum** | -96.320 | 47.017 | 8.005 | 58.081 | 221.137 | 153.751 | 98.776 | 341.96 | 77.665 | |

**Table S6:** Table of z scores of brain effect size correlation comparisons between pairs of granular measures.

| P values | **Writing** | **Throwing** | **Scissors** | **Toothbrush** | **Knife (no fork)** | **Spoon** | **Broom** | **Match** | **Box** | **Hand Sum** |
|---|---|---|---|---|---|---|---|---|---|---|
| **Writing** | | | | | | | | | | |
| **Throwing** | 1.35E-9 | | | | | | | | | |
| **Scissors** | 0 | | | | | | | | | |
| **Toothbrush** | 0 | 0 | 0 | | | | | | | |
| **Knife (no fork)** | 0 | 0 | 0 | 0 | | | | | | |
| **Spoon** | 0 | 0 | 0 | 0 | 0 | | | | | |
| **Broom** | 0 | 0 | 0 | 0 | 0 | 0 | | | | |
| **Match** | 0 | 0 | 0 | 0 | 0 | 0 | 0 | | | |

| | | | | | | | | | |
|---|---|---|---|---|---|---|---|---|---|
| **Box** | 0 | 0 | 0 | 0 | 0 | 0 | 0 | 0 | |
| **Hand Sum** | 0 | 0 | 0 | 0 | 0 | 0 | 0 | 0 | 0 |

**Table S7:** Table of p values of brain effect size correlation comparisons between pairs of granular measures.
